# Mechanism of Action of a Prostaglandin E_2_ Receptor Antagonist with antimicrobial activity against *Staphylococcus aureus*

**DOI:** 10.1101/2020.01.21.914200

**Authors:** Elizabeth A. Lilly, Mélanie A. C. Ikeh, Paul L. Fidel, Mairi C. Noverr

**Author notes:** Address correspondence to Mairi C. Noverr. Department of Molecular and Cell Biology, School of Natural Sciences, University of California, Merced, California, USA. E.A.L. and M.A.C.I. contributed equally to this article as first authors.

## Abstract

Our laboratory recently reported that the EP_4_ receptor antagonist, L-161,982, had direct growth-inhibitory effects on *Staphylococcus aureus in vitro* and in vivo, reducing microbial burden and providing significant protection against lethality in models of *S. aureus* monomicrobial and polymicrobial intra-abdominal infection. This antimicrobial activity was observed with both methicillin-sensitive and methicillin-resistant *S. aureus* (MRSA), as well as other Gram-positive bacteria. The antimicrobial activity of L-161,982 was independent of EP_4_ receptor inhibitory activity. In this study, we investigated the mechanism of action (MOA) of L-161,982, which contains a sulfonamide functional group. However, results demonstrate L-161,982 does not affect folate synthesis (sulfonamide MOA), oxidative stress, or membrane permeability. Instead, our results suggest that the inhibitor works via effects on inhibition of the electron transport chain (ETC). Similar to other ETC inhibitors, L-161,982 exposure results in a small colony size variant phenotype and inhibition of pigmentation, as well as significantly reduced hemolytic activity, and ATP production. In addition, L-161,982 potentiated the antimicrobial activity of another ETC inhibitor and inhibition was partially rescued by supplementation with nutrients required for ETC auxotrophs. Taken together, these findings demonstrate that L-161,982 exerts antimicrobial activity against MRSA via inhibition the ETC, representing a new member of a potentially novel antimicrobial drug class.

## Introduction

*Staphylococcus aureus* is a clinically significant gram positive bacterial pathogen, causing a wide range of nosocomial infections ranging from pneumonia and skin or surgical site infections, to systemic bloodstream infections and sepsis. S. aureus is the leading cause of infection in critically ill and injury patients [1]. There has been an alarming increase in methicillin-resistant *S. aureus* strains (MRSA) and multi-drug resistant (MDR) strains. These resistant strains are prevalent in both hospital-associated and community-acquired infections, which are associated with prolonged hospitalization and treatment costs and mortality rates of 20-50% [2, 3]. Is it estimated that >200,000 MRSA infections occur per year, costing approximately $3.3 billion [4].

MRSA also causes polymicrobial infections, including systemic infections with the fungal pathogen *Candida albicans* [5]. Using animal models, it was demonstrated that polymicrobial infection with *C. albicans* and *S. aureus* causes synergist effects on mortality compared with monomicrobial infections [6–8]. Our laboratory discovered that the immunomodulatory eicosanoid, prostaglandin E2 (PGE_2_), plays a key role in the lethal inflammatory response during polymicrobial intra-abdominal infection (IAI) using a mouse model of infection [6, 9]. In studies designed to uncover key PGE_2_ biosynthesis/signaling components involved in the response, selective eicosanoid enzyme inhibitors and receptor antagonists were selected and pre-screened for antimicrobial activity against *C. albicans* or *S. aureus*. Unexpectedly, we found that the EP_4_ receptor antagonist, L-161,982, had direct growth-inhibitory effects on *S. aureus in vitro* at the physiological concentration required to block PGE_2_ interaction with EP_4_ [10]. This antimicrobial activity was observed with both methicillin-sensitive and methicillin-resistant *S. aureus* (MRSA), with planktonic MIC and MBC values of 50 μg/ml and 100 μg/ml, respectively. In addition, L-161,982 inhibited *S. aureus* biofilm formation and had activity against pre-formed mature biofilms.

In vivo testing demonstrated that treatment of mice with L-161,982 following intra-peritoneal inoculation with a lethal dose of MRSA significantly reduced microbial burden and enhanced survival [9]. Furthermore, L-161,982 protected mice against the synergistic lethality induced by co-infection with *C. albicans* and *S. aureus*. Importantly, administration of L-161,982 alone did not cause any overt signs of toxicity, with no morbidity or mortality observed in control animals. An alternative EP_4_ receptor antagonist exerted no antimicrobial or protective effects, indicating that the antimicrobial activity of L-161,982 is independent of its effects on inhibiting EP_4_ receptor activity. The goal of these studies was to further investigate the antimicrobial mechanism of action (MOA) of L-161,982, testing mechanisms associated with major classes of antimicrobials.

## MATERIALS AND METHODS

### Strains and growth conditions

The methicillin-resistant *S. aureus* strain NRS383 was obtained from the Network on Antimicrobial Resistance in *S. aureus* (NARSA) data bank. As NRS383 has little or no hemolytic activity or pigmentation, *S. aureus* strain 43300 (ATCC) was used for the hemolysis and staphyloxanthin assays. Frozen stocks were obtained at −80°C and streaked onto trypticase soy agar (TSA) plates at 37°C prior to use. A single colony was transferred to 10 ml of trypticase soy broth (TSB) and shaken at 37°C overnight. On the following day, the overnight culture was diluted 1:100 in fresh growth medium and shaken at 37°C for 3 h to obtain cells in mid-log growth phase.

### Effect of exogenous thymidine on antibacterial activity of L-161,982

*S. aureus* was cultured to mid-log phase in cation-adjusted Mueller-Hinton II broth with shaking at 37°C. Cultures were further diluted to yield a final concentration of 5×10^5^ cells/ml. and treated with drugs alone or in combination with 200 µg/ml of thymidine (Sigma). Drugs were L-161,982 (50 µg/ml) or 40 µg/ml of trimethoprim-sulfamethoxazole (SXT) at a ratio of 1:19 (2 µg/ml of trimethoprim and 38 µg/ml of sulfamethoxazole; Sigma) as a positive control. Cultures were incubated at 37°C without shaking and aliquots were removed at t(0), 2, 4, 6, and 24 hours. CFUs were enumerated by plating various dilutions on TSA plates and incubating at 37°C for 24 hours. Results were normalized to negative control cultures containing no drug at each time point.

### Cytoplasmic leakage assay

Cytoplasmic leakage assays were performed as previously described with modifications [11]. Briefly, *S. aureus* was cultured to mid-log phase in TSB with shaking at 37°C. Bacteria were then harvested and resuspended in PBS to a concentration of ∼10^8^/ml. Cultures were treated with L-161,982 (50 µg/ml) or Chlorhexidine (4 µg/ml), a positive control, and incubated at 37°C with shaking. Culture supernatants were monitored over time for cytoplasmic leakage via OD_260_ measurements using a spectrophotometer (Biomate 3S, Thermo Scientific). For this, aliquots were removed at 0.5, 1, 2, and 3 hours and centrifuged for 5 min at 10,000 x *g* at room temperature to remove cellular debris before taking OD_260_ measurements. Negative control cultures containing no drug were used to calculate change in absorbance (ΔA_260_) at each time point.

### Generation of hydroxyl radicals

Antibiotic induced hydroxyl radical production was measured as previously described [12]. Overnight cultures of *S. aureus* were diluted 1:100 in TSB and grown for 3h at 37°C with shaking to mid-log phase. Cultures were diluted in TSB to OD_600_ = 0.05 and L-161,982 was added at a final concentration of 50 μg/ml +/-H_2_O_2_ (1 mM; Sigma) or thiourea (150 mM; Sigma). Cultures were incubated at 37°C with shaking and aliquots were removed at t(0) and at 1 h intervals duplicate, diluted 1:100 and optical density measured at 600 nm (OD_600_) or serially diluted and plated onto SDA and grown at 37°C for 24 h for CFU enumeration.

### Hemolysis Assay

Hemolytic activity was measured using Mueller-Hinton agar plates containing 5% defribinated sheep blood (Thermo Fisher) +/-L-161,982 at a final concentration of 50 μg/ml. To make uniform lawns, *S. aureus* was spread in 1 cm spots and plates were incubated at 37°C for 24 h followed by 24 h incubation at 4°C. Plates were placed on a light box and digitally photographed and images were analyzed using Photoshop CS3. The zone of red blood cell clearance was defined by selecting the area of clearance proximal to the area of growth using the color selection tool set to 25% threshold. The width of each zone was measured at 0, 90, 180, 270° around growth spots (2-3 spots/group) using the ruler tool. Measurements were normalized to petri plate gridlines (13 mm) for each plate.

### Staphyloxanthin Assay

*S. aureus* was diluted 100-fold in 50 ml of TSB containing 50 or 100 µg/ml of L-161,982 and cultured shaking at 37°C for 48 h. Control cultures were grown in broth alone. Cells were harvested by centrifugation (4000 rpm) and washed twice with PBS. Cells from each culture were counted and normalized. Normalized cell suspensions were centrifuged (4000 rpm), resuspended in 1 ml of methanol and incubated at 55°C for 30 minutes to extract pigment. Suspensions were centrifuged at maximum speed (14000 rpm) to remove cell debris, leaving the supernatant containing pigment. Optical densities of supernatants were measured at 465 nm using a spectrophotometer.

### ATP detection assay

*S. aureus* was diluted 100-fold in 10 ml of fresh Mueller Hinton II broth which was used as the inoculum. The inoculum was incubated with various concentrations of L-161,982 (12.5-200 µg/ml) in a 96-well plate in triplicate for 5 h at 37°C. After incubation, cells ± L-161,982 were stained with LIVE/DEAD Baclight bacterial viability stain (Invitrogen) and viable cells were counted. Samples were normalized to viable cell number before transferring to an opaque-walled 96-well plate. An equal volume of BacTiter-Glo™ Reagent (Promega) was added. The plate was allowed to develop for 5 min at room temperature. Luminescence was quantified using a BioTek Synergy 2 plate reader. A media alone control was used to detect background luminescence.

### Electron Transport Chain Inhibitor and Nutritional Rescue Kill Kinetic Assessment

*S. aureus* was cultured to mid-log phase in cation-adjusted Mueller-Hinton II broth with shaking at 37°C. Cultures were further diluted to yield a final concentration of 5×10^5^ cells/ml and treated with each drug alone, L-161,982 (100 µg/ml) or HQNO (10 or 20 µg/ml) (Cayman Chemical), and in various combinations of both for effects on kill kinetics as previously described [13]. *S. aureus* alone containing no drug was used as the positive control. Cultures were incubated at 37°C with shaking and aliquots were removed at t(0), 4, 8, 12, and 24 h. CFUs were enumerated by plating various dilutions on TSA plates and incubating at 37°C for 24 h. Log change in CFU was calculated between each culture at the 24 h endpoint. For rescue studies, *S. aureus* was cultured to mid-log phase in cation-adjusted Mueller-Hinton II broth with shaking at 37°C. Cultures were further diluted to yield a final concentration of 5×10^5^ cells/ml and treated with L-161,982 (100 µg/ml) alone or in combination with 40 mM Pyruvate and 0.1mM uracil. *S. aureus* alone containing no drug was used as the positive control. Cultures were incubated at 37°C with shaking and aliquots were removed at t(0), 4, 8, 12, and 24 h. CFUs were enumerated as above.

### Statistical Analysis

In all assays, samples were analyzed in duplicate and independently repeated. Statistical differences among groups were analyzed using one-way ANOVA and comparisons of each experimental group with control groups were analyzed using the Student’s unpaired t test (2 tailed, unequal variance). In all tests, differences were considered significant at *P* < 0.05. All statistical analyses were performed with Graph Pad Prism software.

## RESULTS

### Growth inhibitory effect of L-161,982 is not due to inhibition of folate synthesis

Examination of the chemical structure of L-161,982 revealed the existence of several functional groups found in different classes of antibiotics. Notably, it has a sulfonamide functional group (-S(=O)_2_-NR_2_), which consists a sulfonyl group connected to an amine group (Fig. 1), similar to the first commercial synthetic antibiotics (sulfonamides or sulfa drugs). The sulfonamide group is located within L-161,982, resulting in a secondary amine group, similar to the sulfonamide antibiotic sulfamethoxazole (SXT). Sulfonamides are bacteriostatic antibiotics that act as competitive inhibitors of the enzyme dihydropteroate synthase (DHPS) involved in folate synthesis, which leads to defective thymidine production and subsequent DNA synthesis [14, 15]. The inhibitory effects of sulfonamides can be antagonized by extracellular thymidine, which provides bacteria an alternative means to folate synthesis [16, 17]. However, in contrast to SXT (positive control drug), addition of exogenous folate did not reverse the growth inhibition exerted by L-161,982 *S. aureus* growth, suggesting the mechanism of action of this drug is unrelated to the sulfonamide group (Fig. 2).

**Figure 1.**
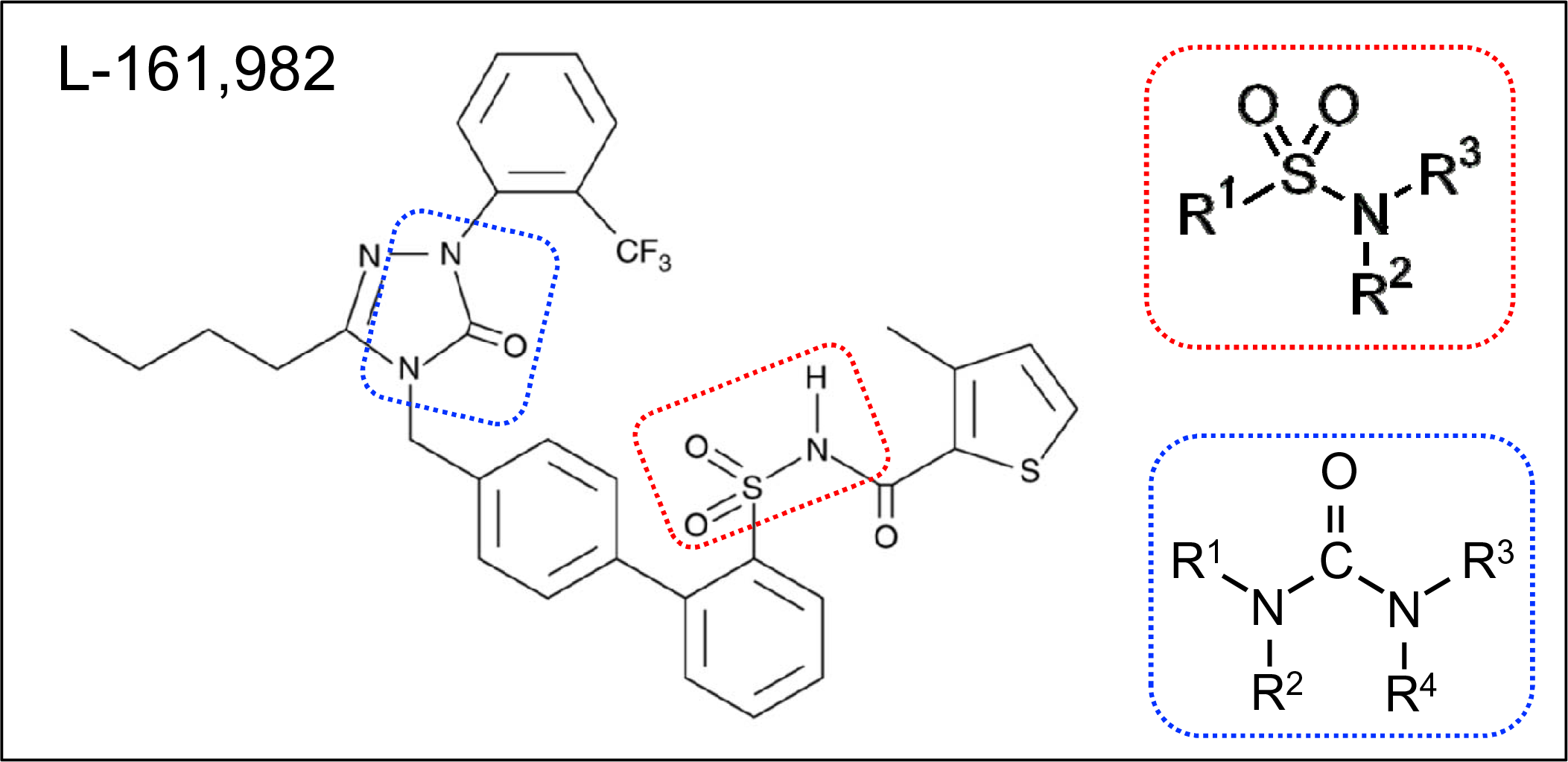
Chemical structure of L-181, 982 N-4’-3-Butyl-1,5-dihydro-5-oxo-1-2-trifluoromethylphenyl-4H-1,2,4-triazol-4-ylmethyl1,1’-biphenyl-2-ylsulfonyl-3-methyl-2-thiophenecarboxamide. Dotted red rectangle outlines the sulfonamide group and dotted blue outline denotes the urea group. Insets shows basic chemical structure of a sulfonamide group and urea group.

**Figure 2.**
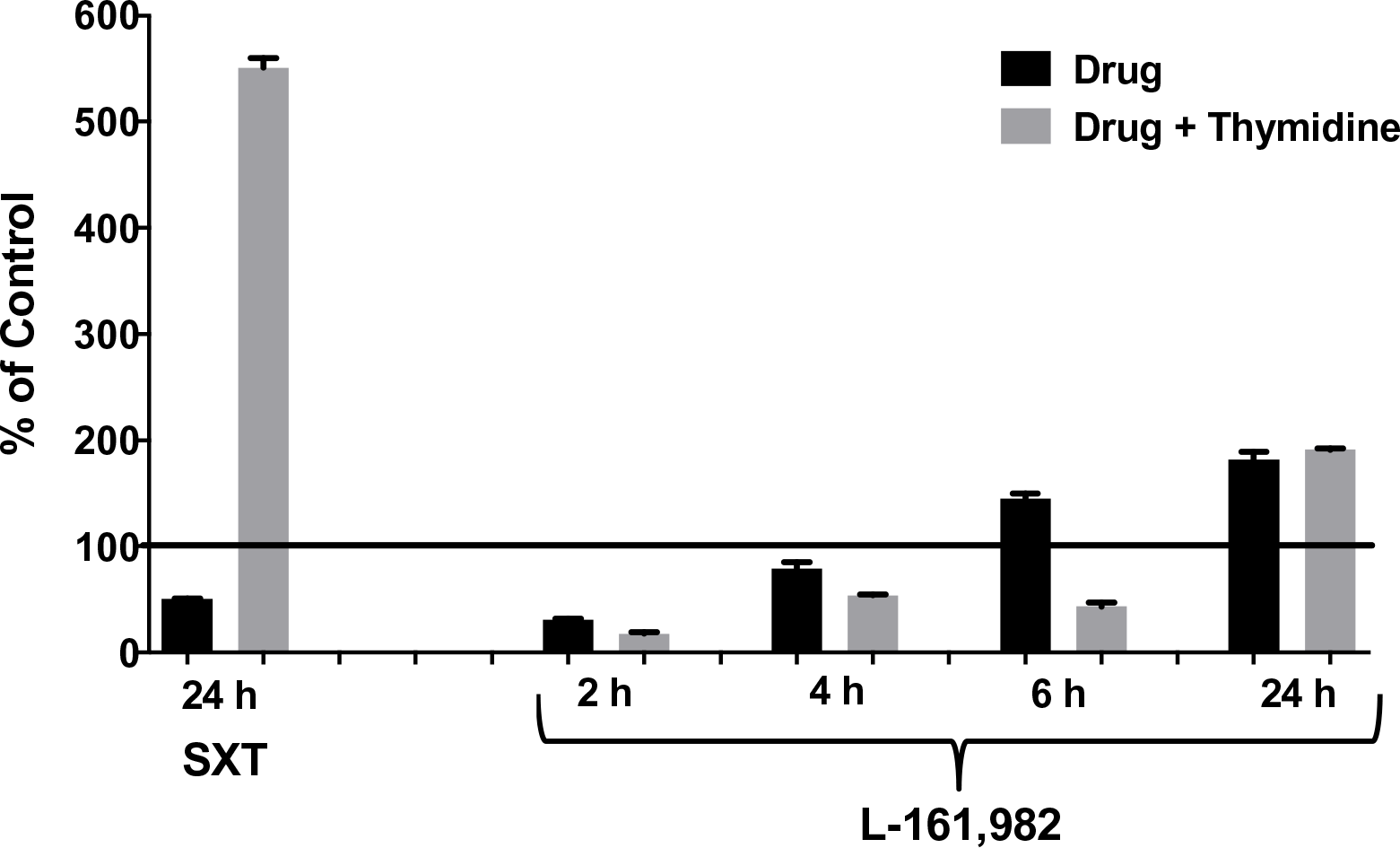
Growth inhibitory effects of L-161,982 are not rescued by exogenous thymidine. *S. aureus* strain NRS383 was grown in cation-adjusted Mueller-Hinton II broth alone (control), with 50 g/ml of L-161,982 or with 40 g/ml of SXT (positive control) ± 200 µg/ml of thymidine. Cultures were incubated at 37°C non-shaking and aliquots were removed at various time points and cultured for growth (CFU). Results are expressed as % of control at each time point. Mean values from two independent experiments are shown.

### L-161,982 does not induce membrane damage or bacteriolysis

Another major target of several classes of antibiotics is the bacterial cell membrane. To monitor membrane damage, the release of nucleic acids and other intracellular 260 nm absorbing material was measured in culture supernatants at 60 min following exposure to drugs and normalized to percent of initial OD. Treatment with L-161,982 did not lead to increases in 260 nm absorbing material (94% ± 1.0), similar to untreated control cultures (90% ± 7.0). However, treatment with a drug known to cause bacteriolysis (chlorhexidine) lead to a significant increase in OD_260_ absorbing material (151.2% ± 9.5; p<0.02 vs untreated control) after 60 min. These findings indicate no loss of bacterial cell membrane integrity occurs following L-161,982 treatment.

### The inhibitory of action of L-161,982 on *S. aureus* growth is not rescued by inhibitors of oxidative stress

A previous study had reported that the inhibitory effect of L-161,982 on the proliferation of skeletal muscle myoblasts was due to its ability to induce production of high levels of intracellular reactive oxygen species (ROS) that could be rescued by co-treatment with the antioxidants, N-acetyl cysteine or sodium ascorbate [18]. Co-treatment with these antioxidants did not circumvent the inhibitory effect of L-161,982 on *S. aureus* growth (Fig. 3A). A common mechanism of cellular death induced by several classes of antibiotics is via stimulation of reactive oxygen species, particularly hydroxyl radicals [12]. Using a potent hydroxyl radical scavenger, thiourea, we further investigated the possibility that L-161,982 induced oxidative stress by generating hydroxyl radicals in *S. aureus*. Again, we found that in the presence of L-161,982, thiourea did not alleviate its inhibitory effect on *S. aureus* growth suggesting the drug did not trigger the generation of hydroxyl radicals (Fig. 3B).

**Figure 3.**
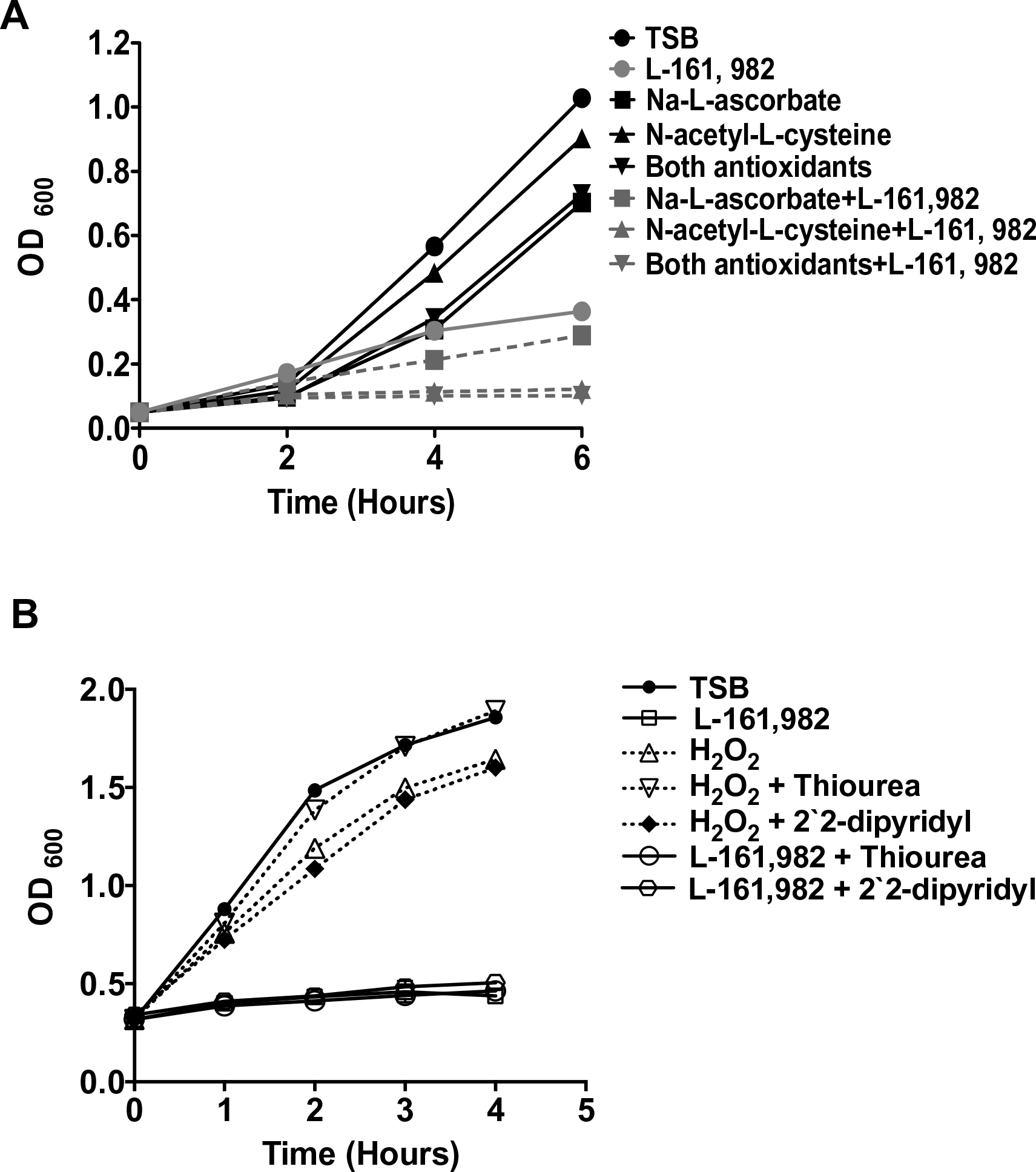
Growth inhibitory effects of L-161,982 are not rescued by inhibitors of oxidative stress. A) Effects of reactive oxygen species inhibitors. Growth of *S. aureus* was monitored for up to 6 h in medium alone or medium supplemented with 15 mM of Na-L-ascorbate, and/or 15mM N-acetyl-L-cysteine, and/or 50 Lg/ml L-161,982. Data shown are representative of two independent experiments. B) Effects of hydroxyl radical scavengers. Growth was monitored for 4 h in medium alone or medium supplemented with 50 g/ml L-161,982 or 1 mM H_2_O_2_ alone, plus 150 mM thiourea, or plus 500 μM 2, 2′ -dipyridyl.

### L-161,982 inhibits activities associated with the electron transport chain including ATP production

In addition to the sulfonamide group, the chemical structure of L 161, 982 contains a urea group connected to two aryl groups (diarylurea) (Fig. 1). Urea derivatives are commonly used as herbicides, interfering with electron transport [19]. Diarylurea compounds have been shown to have similar inhibitory effects on *S. aureus* electron transport, resulting in distinct phenotypes including small colony variant (SCV), inhibition of pigment production, and reduced hemolytic activity [20]. SCVs typically lack a functional electron transport chain and cannot produce virulence factors such as hemolysins or the carotenoid pigment staphyloxanthin [reviewed in [21]]. In the presence of L-161,982, *S. aureus* also grew as small colonies on agar plates (data not shown) and resulted in significant loss of yellow pigmentation (staphyloxanthin) (Fig. 4A & B). Addition of L-161,982 to blood agar resulted in significantly reduced zones of clearance, indicative of reduced hemolysin production (Fig. 5A and B). These defects are all indicative of interruption of electron transport as carotenoid formation as well as toxin production is dependent on an active electron transport [20, 22, 23]. During aerobic respiration, the force required for driving ATP synthesis requires a transmembrane electrical potential (membrane potential), which is generated by the electron transport chain. Addition of L-161,982 to *S. aureus* significantly inhibited ATP production (Fig. 6). Collectively, these findings suggest that the mechanism of action of L-161,982 on *S. aureus* is via inhibition of the electron transport chain.

**Figure 4.**
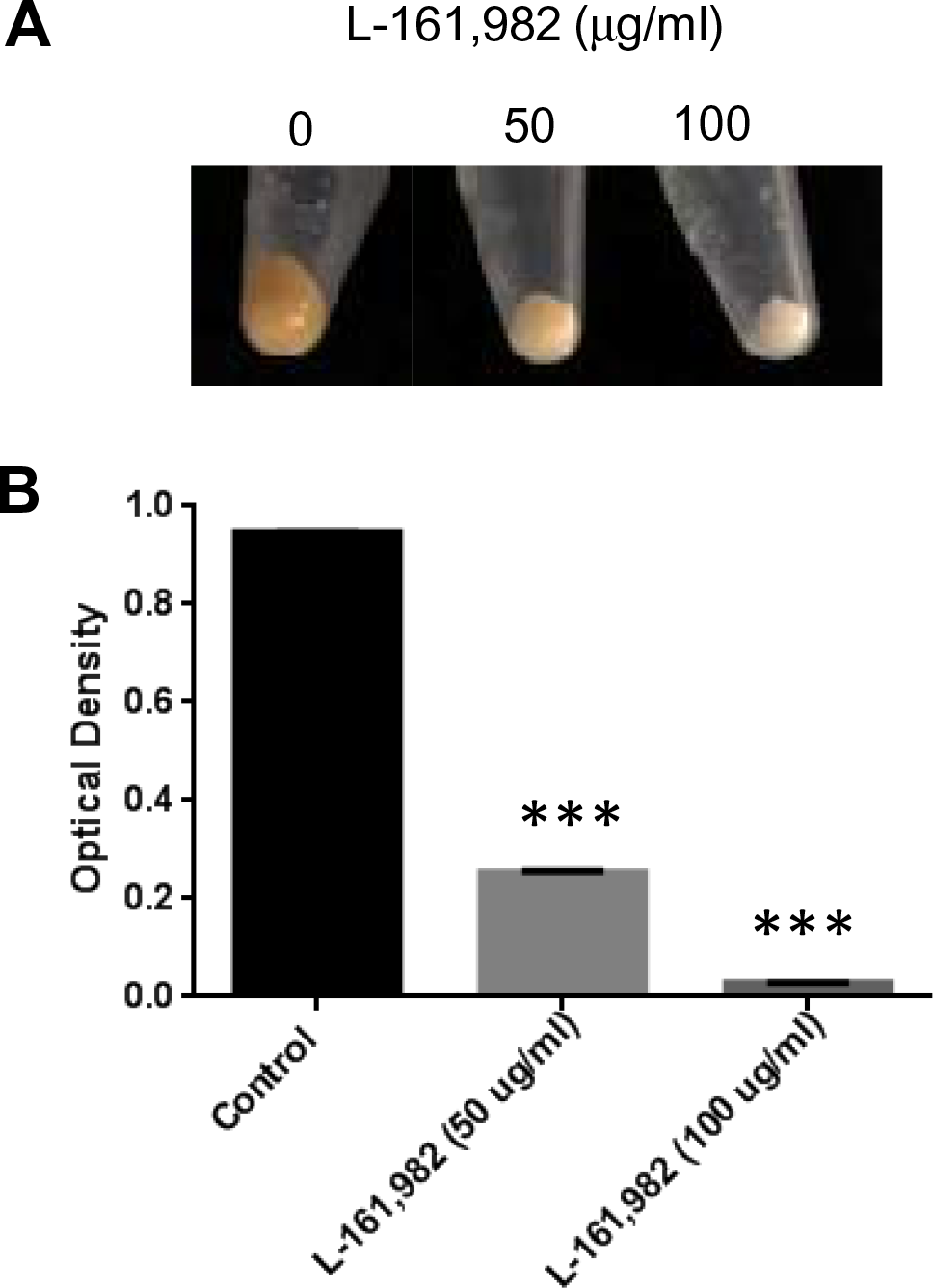
Growth in the presence of L-161,982 inhibits pigment production of S. aureus. Pigment (staphyloxanthin) levels were measured from cultures grown in Mueller-Hinton broth +/-L-161,982 at 37°C for 24h. A) Macroscopic analysis of S. aureus pigment production. B) Pigment was extracted in methanol and optical density measured at 450nm. Data shown are representative of two independent experiments. *** p < 0.001 control vs. drug-treated.

**Figure 5.**
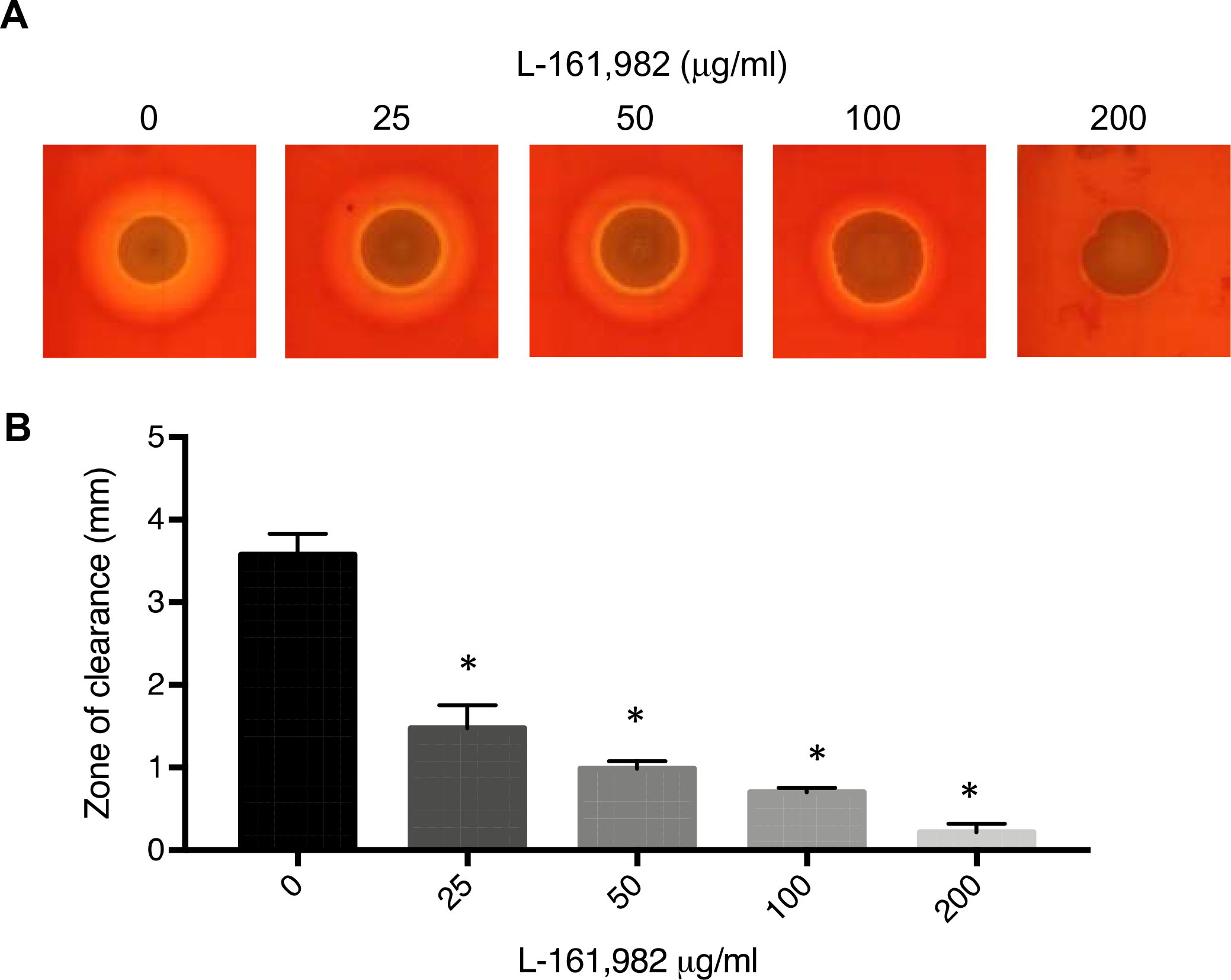
L-161,982 treatment inhibits hemolytic activity of *S. aureus*. A) Macroscopic analysis of S. aureus hemolytic activity. S. aureus was spotted on Mueller-Hinton agar containing sheep blood and/or various concentrations of L-161,982, and incubated at 37C for 24h (0, 25 50, 100) or 72h (200). B) Width of the zone of RBC clearance was measured at 4 points around each S. aureus area of growth in 2 independent replicates. * p < 0.05 control vs. drug-treated.

**Figure 6.**
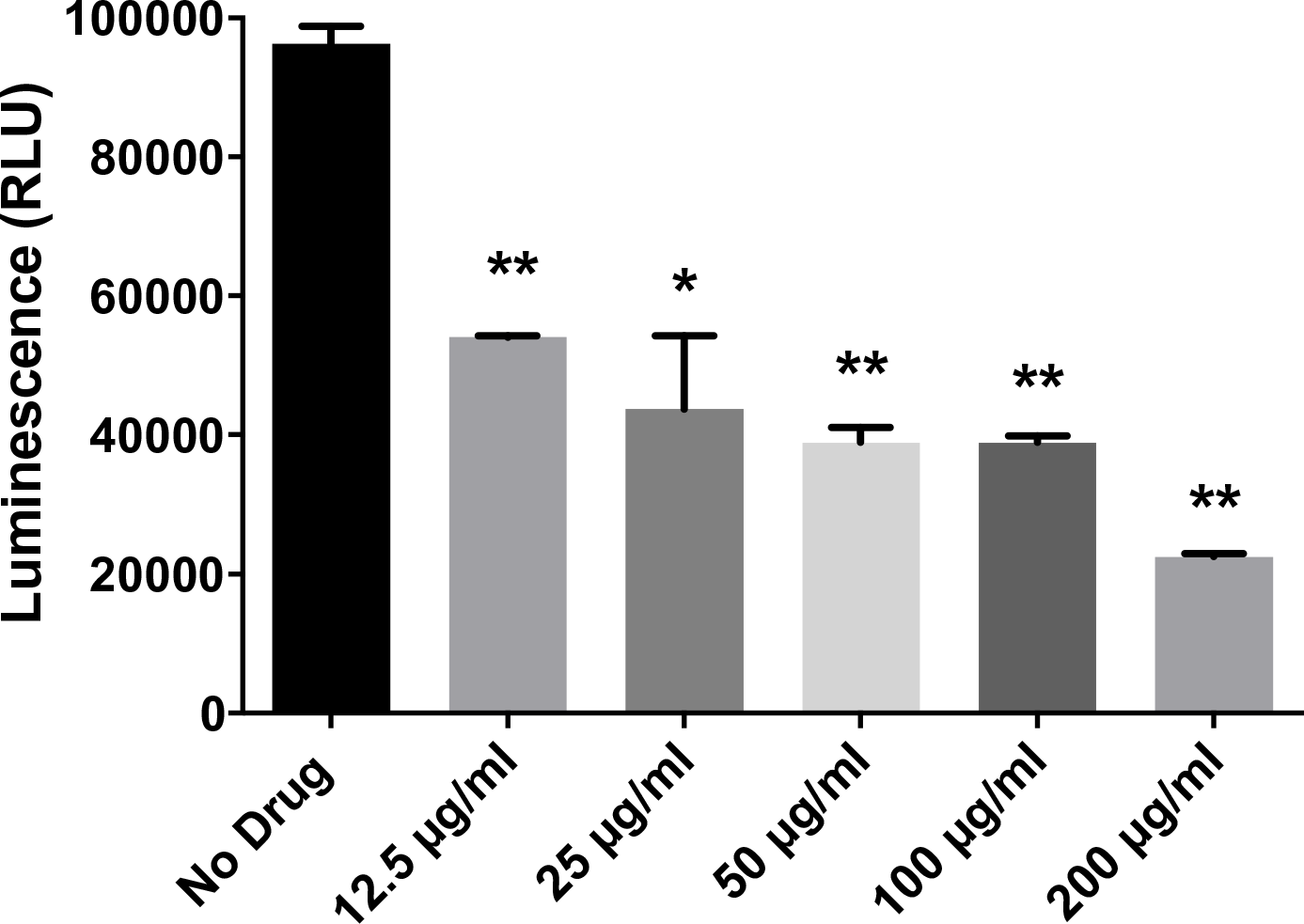
L-161,982 inhibits *S. aureus* ATP production. *S. aureus* was incubated with various concentrations of L-161,982 in a 96-well plate in triplicate for 5 h at 37°C. ATP production was measured in samples normalized to viable cell number using the BacTiter-Glo™ Assay (Promega) and measuring luminescence. Background luminescence from media alone samples was subtracted from experimental samples. Results represent the means +/-standard deviation. Data shown are representative of two independent experiments. ** p < 0.01; *** p < 0.001 control vs drug-treated.

### Inhibitors of *S. aureus* electron transport chain potentiate antibacterial activity of L-161,982, which can be rescued with exogenous uracil/pyruvate

To provide further evidence that L-161,982 targets the electron transport chain and aerobic respiration, we tested for synergy with other electron transport chain inhibitors, including tomatidine (TO), a plant steroidal glycoalkaloid that targets ATP synthase subunit c in *S. aureus* [24] and HQNO (4-hydroxy-2-heptylquinoline-*N*-oxide), a small molecule inhibitor of *S. aureus* electron transport chain produced by *Pseudomonas aeruginosa* [25]. We initially performed standard FIC checkerboard assays; however, neither TO nor HQNO displayed antibacterial activity against our *S. aureus* strain, even at saturating concentrations (data not shown). This is likely due to the fact that these inhibitors exert antibacterial activity against small colony variants of *S. aureus*. Therefore, calculation of FIC indices was not possible. It is important to note that we also observed no synergy with SXT, an antibiotic that targets folic acid synthesis (data not shown). Therefore, we performed kinetic kill assays for assessing the impact of electron transport chain inhibitors in combination with L-161,982 as previously described for combined action of HQNO on TO [13]. Similar to these previous studies with TO, HQNO improved the antibacterial activity of L-161,982 (Fig. 7A). Exposure to L,161,982 or HQNO had a modest effect on the growth of *S. aureus*, leading to a reduction of *S. aureus* counts by ∼0.5 log10 CFU/ml compared to that in the control culture after 24 h. However, in combination these drugs effectively suppressed growth, leading to a significant difference of 2.0 log10 CFU/ml compared to control by 24 h. To rescue the effect of L-161,982 on inhibition of electron transport, media was supplemented with nutrients required for cells lacking a functional respiratory chain including pyruvate and uracil [26]. In *S. aureus*, both pyruvate and uracil are also required for anaerobic growth [27]. Addition of these nutrients improved the growth of *S. aureus* in the presence of L-161,982, resulting in a reproducible increase in growth by 12 h, which was sustained at 24 h (Fig. 7B). Together, these data support confirm that L-161,982 target the electron transport chain and aerobic respiration of *S. aureus*.

**Figure 7.**
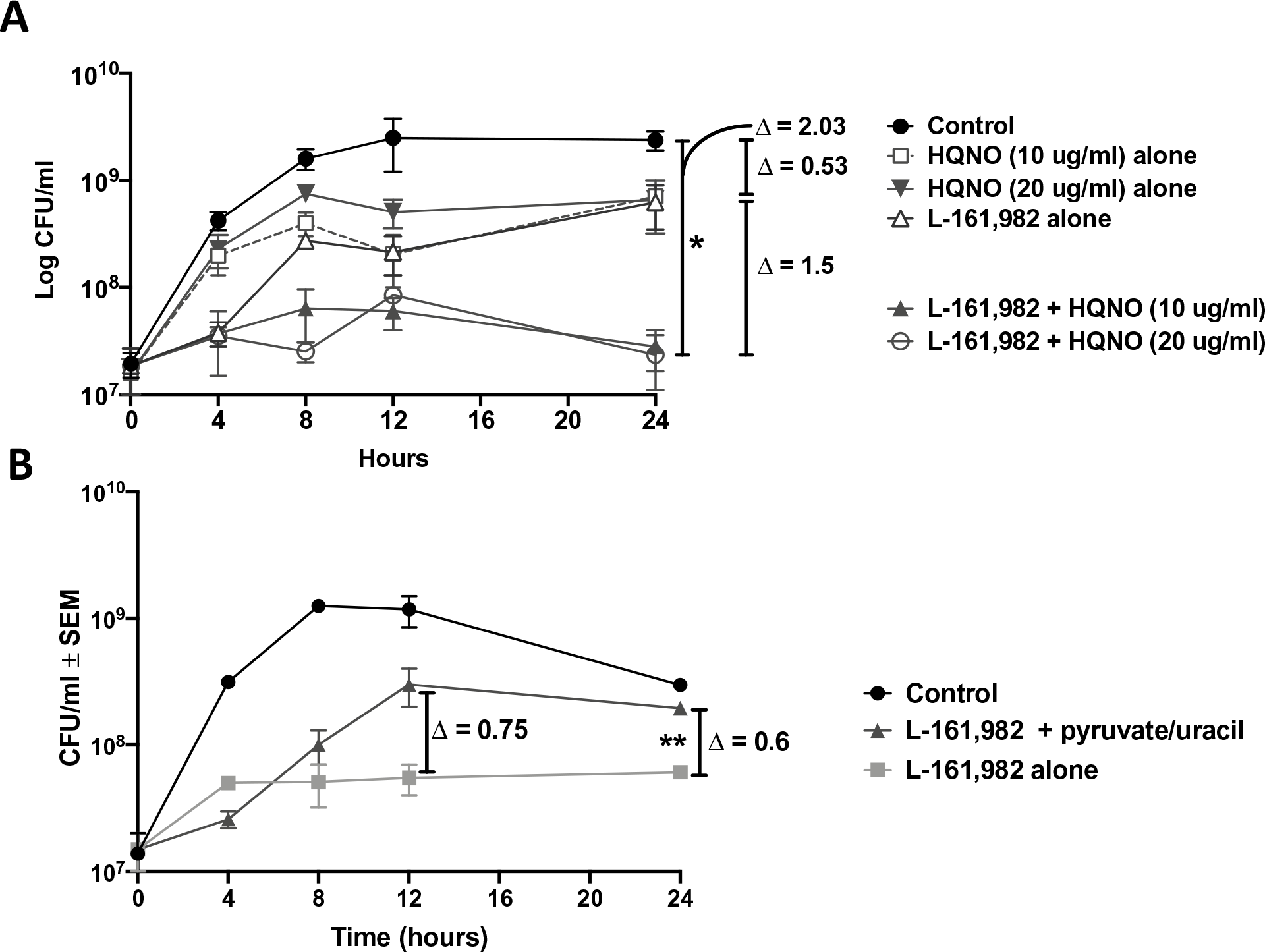
Effect of electron transport chain inhibitors or rescue nutrients on kill kinetics of L-161,982 against *S. aureus*. *S. aureus* was incubated with 100 ug/ml L-161,982 +/-A) 10 or 20 Lg/ml HQNO (electron transport chain inhibitor) or B) 40 mM pyruvate and 0.1 mM uracil (nutrients required during inhibition of electron transport chain and anaerobic respiration). Cultures were incubated at 37°C with shaking and aliquots were removed at t(0), 4, 8, 12, and 24 h. CFUs were enumerated by plating various dilutions on TSA plates and incubating at 37°C for 24 h. Log change in CFU was calculated between groups at 12 or 24 h endpoint. Data shown are representative of two independent experiments. *P<0.05; control vs. L-161,982 +10/20 Lg/ml HQNO. **P<0.05; L-161,982 vs. L-161,982 + pyruvate/uracil.

## DISCUSSION

Antibiotic classification is based upon mechanism of action and drug-target interaction. The three major targets for antibiotics include cell wall/membrane synthesis/integrity, nucleic acid synthesis, and protein synthesis. We recently published that the mammalian EP_4_ receptor antagonist, L-161,982, had direct growth-inhibitory effects on *S. aureus in vitro* at the physiological concentration required to block PGE_2_ interaction with EP_4_ [10]. The antimicrobial activity of L-161,982 was not related to inhibitory effects on host EP_4_ receptor. Therefore, it was unlikely that the mechanism of action of growth inhibition was related to inhibition of any potential receptor homologs in *S. aureus*. The chemical structure of L-161,982 provided some leads on mechanism of action including a sulfonamide group and a several urea groups. Through a series of experiments we also ruled out several potential mechanisms of action including inhibition of folate synthesis and thymidine production, generation of oxidative stress, and membrane damage.

In *S. aureus*, it was previously reported that several diaryl urea compounds exert antibacterial activity via inhibition of the electron transport chain [20]. L-161,982 also contains a urea group flanked by two aryl groups, and may have similar effects on *S. aureus*. The electron transport chain enables aerobic respiration and subsequent ATP production and is a less common target for antimicrobial agents. Distinct phenotypic characteristics associated with interruptions in electron transport include slow growth as cell wall biosynthesis is dependent on copious amounts of ATP, decreased pigmentation as carotenoid formation (staphyloxanthin) requires electron transport, and production of hemolysins [20, 28]. Our initial observation that L-161,982 slowed growth of *S. aureus* suggested a potential effect on the electron transport chain [10]. In agreement with this observation, exposure to L-161,982 also resulted in small colony variant (SCV) phenotype similar to diaryl urea compounds [20]. Subsequent testing revealed that L-161,982 inhibited pigment production and hemolytic activity, also consistent with inhibition of the electron transport chain. L-161,982 inhibited ATP as the end product of the electron transport chain, which served as the most direct evidence for its mode of action. Another ETC inhibitor, HQNO, also potentiated the antibacterial effect of L-161,982, similar to studies with HQNO and tomatidine, which targets *S. aureus* ATP synthase subunit C [13, 24]. Further evidence was provided by partial rescue of growth inhibition by addition of exogenous pyruvate and uracil, which are required for growth of respiratory chain auxotrophs and anaerobic growth of *S. aureus* [26, 27].

In summary, results from these studies support the concept that the mechanism of action of L-161,982 is via inhibition of the electron transport chain and subsequent ATP production. Further studies are required to establish the specific step or component of the electron transport chain inhibited by L-161,982. This information would be important in determining species specificity of the compound. There is the possibility that L, 161, 982 may also exhibit antagonism with other classes of antibiotics. For example, *S. aureus* SCVs are perhaps better known as clinical isolates that display resistance to aminoglycosides, antibiotics that target the ribosomal 30S subunit and inhibit protein synthesis. Aminoglycosides require ATP for uptake by *S. aureus*, therefore inhibition of respiration is associated with drug resistance [28]. For this reason, L-161,982 may exert antagonism with this class of antibiotic. But with that withstanding, L-161,982 is very effective against MRSA, both in vitro and in vivo [10]. This is due to the fact that L, 161, 982 targets a completely different biological function than methicillin, which targets cell wall synthesis. Therefore, further development of electron transport chain inhibitors could represent a clinically important strategy to treat MRSA infections, particularly considering administration of L-161,982 dramatically improved survival during both systemic *S. aureus* infection and polymicrobial sepsis induced by *S. aureus* and *C. albicans* [10].

## REFERENCES

1. Sampedro GR, Bubeck Wardenburg J. Staphylococcus aureus in the Intensive Care Unit: Are These Golden Grapes Ripe for a New Approach? J Infect Dis. 2017;215(suppl_1):S64–S70. doi: 10.1093/infdis/jiw581. PubMed PMID: 28003353.

2. Dantes R, Mu Y, Belflower R, Aragon D, Dumyati G, Harrison LH, et al. National burden of invasive methicillin-resistant Staphylococcus aureus infections, United States, 2011. JAMA Intern Med. 2013;173(21):1970–8. doi: 10.1001/jamainternmed.2013.10423. PubMed PMID: 24043270.

3. Calfee DP. Trends in Community Versus Health Care-Acquired Methicillin-Resistant Staphylococcus aureus Infections. Curr Infect Dis Rep. 2017;19(12):48. Epub 2017/11/03. doi: 10.1007/s11908-017-0605-6. PubMed PMID: 29101576.

4. Gidengil CA, Gay C, Huang SS, Platt R, Yokoe D, Lee GM. Cost-effectiveness of strategies to prevent methicillin-resistant Staphylococcus aureus transmission and infection in an intensive care unit. Infect Control Hosp Epidemiol. 2015;36(1):17–27. doi: 10.1017/ice.2014.12. PubMed PMID: 25627757; PubMed Central PMCID: PMCPMC4311265.

5. Nair N, Biswas R, Götz F, Biswas L. Impact of Staphylococcus aureus on pathogenesis in polymicrobial infections. Infection and immunity. 2014;82(6):2162–9. Epub 2014/03/18. doi: 10.1128/IAI.00059-14. PubMed PMID: 24643542; PubMed Central PMCID: PMCPMC4019155.

6. Peters BM, Noverr MC. Candida albicans-Staphylococcus aureus polymicrobial peritonitis modulates host innate immunity. Infect Immun. 2013;81(6):2178–89. Epub 2013/04/03. doi:10.1128/iai.00265-13. PubMed PMID: 23545303; PubMed Central PMCID: PMC3676024.

7. Nash EE, Peters BM, Palmer GE, Fidel PL, Noverr MC. Morphogenesis is not required for Candida albicans-Staphylococcus aureus intra-abdominal infection-mediated dissemination and lethal sepsis. Infect Immun. 2014;82(8):3426–35. doi: 10.1128/iai.01746-14. PubMed PMID: 24891104; PubMed Central PMCID: PMC4136217.

8. Nash EE, Peters BM, Fidel PL, Noverr MC. Morphology-Independent Virulence of Candida Species during Polymicrobial Intra-abdominal Infections with Staphylococcus aureus. Infect Immun. 2015;84(1):90–8. doi: 10.1128/iai.01059-15. PubMed PMID: 26483410; PubMed Central PMCID: PMC4694008.

9. Ikeh MAC, Fidel PL, Noverr MC. Identification of Specific Components of the Eicosanoid Biosynthetic and Signaling Pathway Involved in pathological inflammation during Intra-abdominal Infection with. Infection and immunity. 2018. Epub 2018/05/07. doi: 10.1128/IAI.00144-18. PubMed PMID: 29735520.

10. Ikeh MAC, Fidel PL, Jr., Noverr MC. Prostaglandin E2 Receptor Antagonist with Antimicrobial Activity against Methicillin-Resistant Staphylococcus aureus. Antimicrobial agents and chemotherapy. 2018;62(3). Epub 2017/12/22. doi: 10.1128/AAC.01920-17. PubMed PMID: 29263068; PubMed Central PMCID: PMC5826135.

11. Carson CF, Mee BJ, Riley TV. Mechanism of action of Melaleuca alternifolia (tea tree) oil on Staphylococcus aureus determined by time-kill, lysis, leakage, and salt tolerance assays and electron microscopy. Antimicrobial agents and chemotherapy. 2002;46(6):1914–20. PubMed PMID: 12019108; PubMed Central PMCID: PMCPMC127210.

12. Kohanski MA, Dwyer DJ, Hayete B, Lawrence CA, Collins JJ. A common mechanism of cellular death induced by bactericidal antibiotics. Cell. 2007;130(5):797–810. doi: 10.1016/j.cell.2007.06.049. PubMed PMID: 17803904.

13. Boulanger S, Mitchell G, Bouarab K, Marsault E, Cantin A, Frost EH, et al. Bactericidal Effect of Tomatidine-Tobramycin Combination against Methicillin-Resistant Staphylococcus aureus and Pseudomonas aeruginosa Is Enhanced by Interspecific Small-Molecule Interactions. Antimicrobial agents and chemotherapy. 2015;59(12):7458–64. Epub 2015/09/24. doi: 10.1128/AAC.01711-15. PubMed PMID: 26392496; PubMed Central PMCID: PMC4649251.

14. Hitchings GH, Burchall JJ. Inhibition of folate biosynthesis and function as a basis for chemotherapy. Adv Enzymol Relat Areas Mol Biol. 1965;27:417–68. PubMed PMID: 4387360.

15. Chernyshev A, Fleischmann T, Kohen A. Thymidyl biosynthesis enzymes as antibiotic targets. Appl Microbiol Biotechnol. 2007;74(2):282–9. Epub 2007/01/11. doi: 10.1007/s00253-006-0763-1. PubMed PMID: 17216455.

16. Zander J, Besier S, Saum SH, Dehghani F, Loitsch S, Brade V, et al. Influence of dTMP on the phenotypic appearance and intracellular persistence of Staphylococcus aureus. Infection and immunity. 2008;76(4):1333–9. Epub 2007/12/26. doi:10.1128/IAI.01075-07. PubMed PMID: 18160477; PubMed Central PMCID: PMCPMC2292857.

17. Proctor RA. Role of folate antagonists in the treatment of methicillin-resistant Staphylococcus aureus infection. Clinical infectious diseases: an official publication of the Infectious Diseases Society of America. 2008;46(4):584–93. doi: 10.1086/525536. PubMed PMID: 18197761.

18. Mo C, Zhao R, Vallejo J, Igwe O, Bonewald L, Wetmore L, et al. Prostaglandin E2 promotes proliferation of skeletal muscle myoblasts via EP4 receptor activation. Cell Cycle. 2015;14(10):1507–16. doi: 10.1080/15384101.2015.1026520. PubMed PMID: 25785867; PubMed Central PMCID: PMCPMC4615122.

19. Hosler JP, Yocum CF. Regulation of Cyclic Photophosphorylation during Ferredoxin-Mediated Electron Transport: Effect of DCMU and the NADPH/NADP Ratio. Plant Physiol. 1987;83(4):965–9. PubMed PMID: 16665372; PubMed Central PMCID: PMCPMC1056483.

20. Proctor RA, Dalal SC, Kahl B, Brar D, Peters G, Nichols WW. Two diarylurea electron transport inhibitors reduce Staphylococcus aureus hemolytic activity and protect cultured endothelial cells from lysis. Antimicrobial agents and chemotherapy. 2002;46(8):2333–6. PubMed PMID: 12121901; PubMed Central PMCID: PMCPMC127355.

21. Proctor RA, Kahl B, von Eiff C, Vaudaux PE, Lew DP, Peters G. Staphylococcal small colony variants have novel mechanisms for antibiotic resistance. Clinical infectious diseases: an official publication of the Infectious Diseases Society of America. 1998;27 Suppl 1:S68–74. PubMed PMID: 9710673.

22. Balwit JM, van Langevelde P, Vann JM, Proctor RA. Gentamicin-resistant menadione and hemin auxotrophic Staphylococcus aureus persist within cultured endothelial cells. The Journal of infectious diseases. 1994;170(4):1033–7. PubMed PMID: 7930701.

23. Proctor RA, Balwit JM, Vesga O. Variant subpopulations of Staphylococcus aureus as cause of persistent and recurrent infections. Infect Agents Dis. 1994;3(6):302–12. PubMed PMID: 7889317.

24. Lamontagne Boulet M, Isabelle C, Guay I, Brouillette E, Langlois JP, Jacques PE, et al. Tomatidine Is a Lead Antibiotic Molecule That Targets Staphylococcus aureus ATP Synthase Subunit C. Antimicrobial agents and chemotherapy. 2018;62(6). Epub 2018/04/04. doi: 10.1128/AAC.02197-17. PubMed PMID: 29610201; PubMed Central PMCID: PMC5971568.

25. Mitchell G, Seguin DL, Asselin AE, Deziel E, Cantin AM, Frost EH, et al. Staphylococcus aureus sigma B-dependent emergence of small-colony variants and biofilm production following exposure to Pseudomonas aeruginosa 4-hydroxy-2-heptylquinoline-N-oxide. BMC microbiology. 2010;10:33. Epub 2010/02/02. doi: 10.1186/1471-2180-10-33. PubMed PMID: 20113519; PubMed Central PMCID: PMC2824698.

26. Sullivan LB, Gui DY, Hosios AM, Bush LN, Freinkman E, Vander Heiden MG. Supporting Aspartate Biosynthesis Is an Essential Function of Respiration in Proliferating Cells. Cell. 2015;162(3):552–63. Epub 2015/08/02. doi: 10.1016/j.cell.2015.07.017. PubMed PMID: 26232225; PubMed Central PMCID: PMC4522278.

27. Evans JB. Uracil and pyruvate requirements of anaerobic growth of staphylococci. Journal of clinical microbiology. 1975;2(1):14–7. Epub 1975/07/01. PubMed PMID: 1225927; PubMed Central PMCID: PMC274119.

28. von Eiff C, Peters G, Becker K. The small colony variant (SCV) concept -- the role of staphylococcal SCVs in persistent infections. Injury. 2006;37 Suppl 2:S26–33. doi: 10.1016/j.injury.2006.04.006. PubMed PMID: 16651068.

